# A new cellular platform for studying autophagy

**DOI:** 10.1101/2024.03.28.587157

**Authors:** Korina Goldin-Azulay, Milana Fraiberg, Olena Trofimyuk, Yishai Levin, Nina Reuven, Ekaterina Kopitman, Zvulun Elazar

**Author notes:** Corresponding author - Zvulun Elazar, Department of Biomolecular Sciences The Weizmann Institute of Science 76100 Rehovot, Israel.

## Abstract

Atg8 proteins play a crucial role in autophagy. There is a single Atg8 isoform in yeast, while mammals have up to seven homologs categorized into LC3s and GABARAPs. The GABARAP subfamily consists of GABARAP, GABARAPL1, and GABARAPL2/GATE16, implicated in various stages along the pathway. However, the intricacies among GABARAP proteins are complex and require a more precise delineation.

Here, we introduce a new cellular platform to study autophagy using CRISPR/Cas9-mediated tagging of endogenous genes of the GABARAP subfamily with different fluorescent proteins. This platform allows robust examination of autophagy by flow cytometry of cell populations and monitoring of GABARAP homologs at single-cell resolution using fluorescence microscopy. Strikingly, the simultaneous labeling of the different endogenous GABARAPs allows the identification and isolation of autophagosomes differentially marked by these proteins. Using this system, we found that the different GABARAPs are associated with different autophagosomes. We argue that this new cellular platform will be crucial in studying the unique roles of individual GABARAP proteins in autophagy and other putative cellular processes.

## Introduction

Autophagy, a lysosome-dependent degradation pathway in eukaryotes, plays a critical role in maintaining cellular homeostasis by managing intracellular quality control and enabling cellular adaptation to environmental changes. Dysregulation of autophagy plays a role in aging pathologies, including neurodegeneration, cancer, and inflammation [1]. The Atg8 family of ubiquitin-like proteins plays a fundamental role in autophagy and other processes related to vesicle transport and fusion with the lysosome [1, 2]. The mammalian Atg8 family comprises the LC3 and GABARAP subfamilies, each containing at least three homologs [1–3]. Eliminating all three GABARAPs impairs autophagic flux and fusion with the lysosome [2, 4–7].

In addition to their roles in nonselective autophagy, GABARAPs play a critical role in several types of selective autophagy, including mitophagy, ER-phagy, Golgi-phagy, and centrosome-phagy [8, 9]. Recent studies revealed that GABARAPs also possess non-autophagic functions in immune and inflammatory responses, receptor endocytosis, vesicle trafficking, neurotransmitter transport, cilia formation, and cell adhesion [2, 10]. This suggests functional divergence among different GABARAP isoforms within autophagy and other cellular pathways [5, 11–13]. The main contemporary challenge in studying the individual roles of GABARAPs results is the absence of tools for direct monitoring of distinct isoforms expressed at their physiological levels *in vivo* upon fusion to functional protein tags such as epitopes and fluorescent proteins [14].

Here, we introduce a novel platform for investigating autophagy and related processes involving GABARAPs. Utilizing CRISPR/Cas9 technology, we engineered reporter cell lines with N’-terminal 3xFlag-epitope, fused to bright fluorescent proteins Clover or mScarletI, integrated at their native chromosomal loci. This platform allows *in vivo* characterization of endogenous GABARAPs, individually or in pairs. We demonstrate the amenability of this system to biochemical assays and isolation of autophagosome subsets specifically labeled by various GABARAP homologs. These single or duplex-tagged GABARAPs reporter cells are also primed for high-throughput screening studies by flow cytometry, owing to their robust response to autophagy-inducing environmental conditions.

## Results

### Generation of single and duplex GABARAPs reporter cell lines

Using CRISPR/Cas9, we created single-tagged and double-tagged (duplex) GABARAPs reporter cells. In single tagged reporter cells, GABARAP, GABARAPL1 or GABARAPL2/GATE16 (GATE16 hereafter) were labeled at their N’-terminus with 3xFlag epitope fused to a Clover fluorescent protein [15] at their native chromosomal locations and under their endogenous promotor (*Figure 1A*). Duplex reporter cells were generated from single ^3xFlag-Clover^GABARAP or ^3xFlag-Clover^GATE16 cells by tagging GABARAPL1 or GATE16 (*Figure 1E*) with 3xFlag fused to mScarletI. The latter is a bright red, rapidly maturing fluorescent protein known for its moderate sensitivity to acidic environments [16], including lysosomes, which makes it suitable for autophagic studies. After selection by fluorescence-activated cell sorting (FACS) and genomic sequencing, both reporter cell types were validated for genomic tagging of GABARAPs by western blot analysis (*Figure 1C and 1G*). GABARAP isoform pairs in duplex cells were distinguished by immunoblot analysis, where the mScarletI-tagged isoform migrated slower than the Clover-tagged isoform (*Figure 1G*). Gene-specific editing was also confirmed by knockdown of endogenous GABARAP proteins in both reporter cell types (*Figure 1 – Figure Supplement 1A and Figure Supplement 1B*). In addition, the fluorescence of both reporter cell types was examined and confirmed by confocal microscopy (*Figure 1B and 1F*) and flow cytometry (*Figure 1D and 1H*).

**Figure 1.**
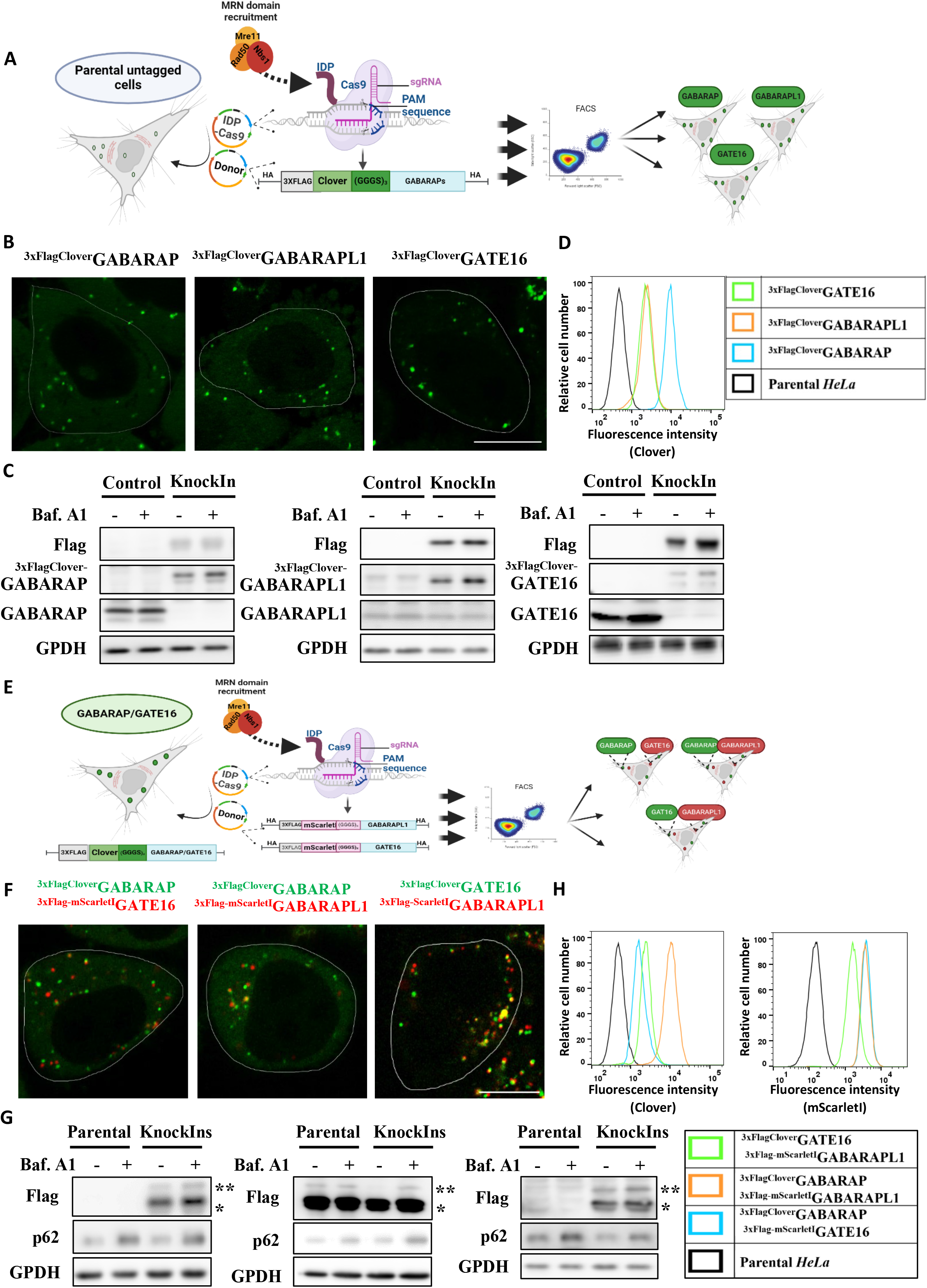
Generation of single and duplex GABARAPs reporter cell lines. **A.** Schematic model of single color GABARAPs knock-in a process mediated by CRISPR/Cas9. SpCas9 was fused at its N-terminus to the intrinsically disordered viral protein (IDP), enhancing the recruitment of the MRN complex. The construct includes a 3xFlag affinity tag followed by Clover fluorescent protein, flanked by 1000bp homology arms (HA). Figure created with BioRender.com **B.** Visualization of endogenously tagged single GABARAPs under basal conditions by super-resolution. Scale bar 10µm. **C.** Endo-tagged GABARAPs expression validated in all reporter cell lines by western blot analysis. The parental *HeLa* cells (control) and single endo tagged GABARAPs reporter cell lines were grown to confluence in a complete medium and treated (where indicated) for the last four h with 0.1 µM Bafilomycin A1. Total protein extracts were analyzed for Flag, GABARAP or GATE16 or GABARAPL1, and GPDH (loading control). **D.** Validation of Clover fluorescence in all single endo tagged GABARAPs reporter cell lines versus parental *HeLa* cells by Flow cytometry under basal conditions. **E.** Schematic model of dual-color GABARAPs knockIn process mediated by CRISPR/Cas9. SpCas9 was fused at its N-terminus to the intrinsically disordered viral protein (IDP), enhancing the recruitment of the MRN complex. The single-color reporter cells expressing GABARAP or GATE16 tagged by 3xFlag affinity tag followed by Clover fluorescent protein were endo-tagged with 3xFlag affinity tag followed by mScarletI fluorescent protein on the other GABARAP family member, producing duplex of endo-tagged pairs of GABARAP proteins. Figure created with BioRender.com **F.** Visualization of endogenously tagged duplex GABARAPs under basal conditions by super-resolution microscopy confocal microscopy. Scale bar 10µm. **G.** Dual-color endo-tagged GABARAPs expression validated in three duplex reporter cell lines by western blot analysis. The parental *HeLa* cells (control) and duplex endo-tagged GABARAPs reporter cell lines were grown to confluence in a complete medium and treated (where indicated) for the last 4 h with 0.1 µM Bafilomycin A1. Total protein extracts were analyzed for Flag, p62, and GPDH (loading control). For Flag protein *-3xFlagClover **-3xFlag-mScarletI **H.** Validation of Clover and mScarletI fluorescence in all duplex endo tagged GABARAPs reporter cell lines versus parental *HeLa* cells by Flow cytometry under basal conditions.

### Analyzing autophagy by single-tagged GABARAP reporter cells

To examine the reliability and suitability of our single-tagged reporter cells for GABARAPs-related studies, cells were exposed to nutrient deprivation to induce autophagy. The endo-tagged GABARAPs response to lysosomal inhibitor Bafilomycin (Baf.A1) and starvation conditions was further examined by immunoblotting (*Figure 2C and Figure 2 – Figure supplement 1A*) and by confocal microscopy (*Figure 2A*). As Depicted in *Figure 2B*, ^3xFlag-Clover^GABARAP, ^3xFlag-Clover^GABARAPL1, and ^3xFlag-Clover^GATE16 were accumulated in response to treatment with Baf.A1 under both basal and starvation conditions. However, this accumulation did not occur with the additional knockout of ATG14, a crucial factor for autophagosome formation (*Figure 2 – Figure supplement 1C*) [17]. The autophagy receptor SQSTM1/p62 was utilized as a positive control and exhibited changes in protein levels consistent with those of endo-tagged GABARAPs (*Figure 2C, Figure 2 – Figure supplement 1A*). Visualization of individual ^3xFlag-Clover^GABARAPs demonstrated their subcellular colocalization with p62 and the lysosomal marker LAMP1 under bulk and starvation conditions – both enhanced upon treatment with Baf.A1 (*Figure 2 – Figure Supplement 2A and Figure Supplement 2B, respectively*). To validate the system compatibility with high-throughput studies, we employed a flow cytometry to quantify endo-tagged GABARAPs response to starvation-induced autophagy [18] and observed a marked change in fluorescence upon induction of autophagy, which was abolished upon knockout of ATG14 (*Figure 2C*).

**Figure 2.**
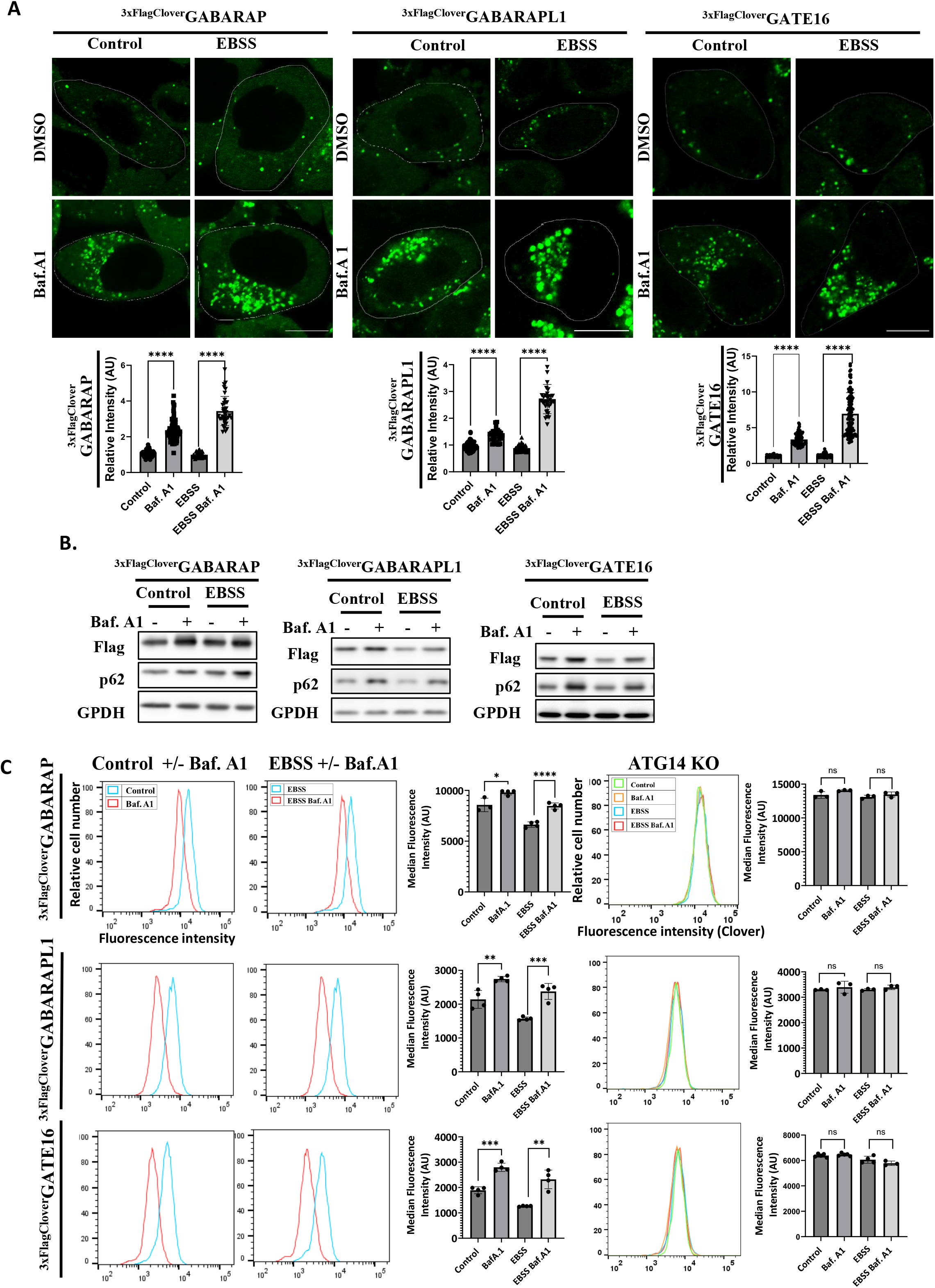
Analyzing autophagy by single-tagged GABARAP reporter cells. **A.** Confocal super-resolution microscopy visualization of single endo tagged GABARAPs under basal and starvation conditions. Cells were incubated in EBSS for 4 hours and treated with 0.1 μM lysosomal inhibitor Bafilomycin A1 for 4 hours (where indicated). Scale bar: 10µm. For analysis, the visualization been done using spinning disk confocal. Relative intensity was calculated using ImageJ. Images were subjected to *maximum projection* and background subtraction using the *rolling ball* function, quantified using ROIs for single cells, and normalized to control. Data are presented with the SEM from three independent experiments, with statistical significance determined by a t-test (****p < 0.0001). **B.** Single endo-tagged GABARAPs response to autophagy-inducing conditions indicated by western blot analysis. Cells were grown to confluence in a complete (control) and starvation (EBSS) medium and treated (where indicated) for the last 4 h with 0.1 µM Bafilomycin A1. Total protein extracts were probed for Flag, SQSTM1 (p62), and GPDH (loading control). **C.** Flow cytometry analysis (FACS) of the Clover fluorescence response in single endo-tagged GABARAPs, with or without Atg14 knockout, to autophagy-inducing conditions. Median Fluorescence Intensity (arbitrary units) was measured in fluorescence units and presented with the SEM from three independent experiments. Statistical significance was determined by a t-test, with **p < 0.01, ***p < 0.001, ****p < 0.0001, ns – insignificant.

These observations confirm the reliability of our single endo-tagged GABARAPs for reporting induction of autophagy or inhibition in autophagic consumption by the lysosome, thus supporting their proper biological functionality and relevance for individual GABARAP-related studies.

### Measuring autophagy with a duplex GABARAPs reporter system

Next, we assayed our GABARAPs duplex platform by applying autophagy-inducing conditions. First, we examined protein levels of endo-tagged GABARAP pairs and, as expected, obtained similar accumulation in all three pairs of GABARAPs, also consistent with p62 accumulation upon Baf.A1 treatment under basal and starvation conditions (*Figure 3 – Figure Supplement 1A*). Confocal microscopy visualization of endo-tagged GABARAP pairs showed a Baf.A1-dependent increase in fluorescent intensity under basal and starvation conditions, in line with western blot (*Figure 3A, Figure 3 - figure supplement 1A, respectively*) and flow cytometry findings (*Figure 3B*). Intriguingly, GABARAP homologs predominantly exhibited distinct vesicular localizations in steady-state conditions, as duplex cell lines revealed only 20-40% overlap between GABARAP protein subfamily members under basal conditions (*Figure 3 - Figure Supplement 1B*). All GABARAP pairs showed colocalization to p62 and Lamp-iRPF670 under basal and starvation conditions that considerably increased upon Baf.A1 treatment (*Figure 3, Supplement 2 and Figure 3, Supplement 3 respectively*). Flow cytometry analysis of the endo-tagged GABARAP pairs showed profound change upon autophagy induction, consistent with findings from western blot and confocal microscopy (*Figure 3B*).

**Figure 3.**
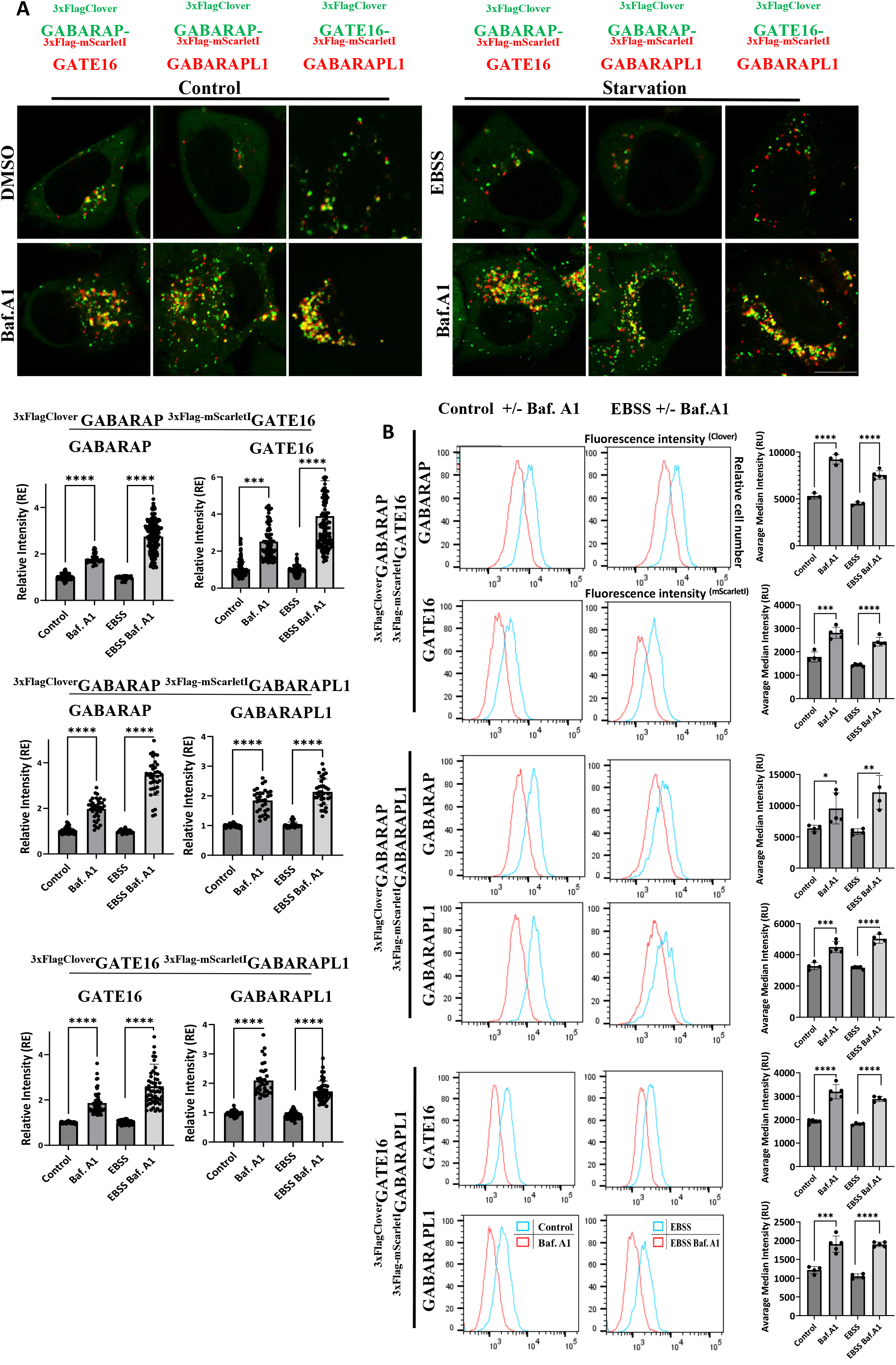
Measuring autophagy with the duplex GABARAPs reporter system. **A.** Confocal super-resolution microscopy representative images of duplex endo tagged GABARAPs under basal and starvation conditions. Cells were incubated in EBSS for 4 hours and treated with 0.1 µM lysosomal inhibitor Bafilomycin A1 for 4 hours (where indicated). Scale bar: 10 µm. For analysis, the visualization been done using spinning disk confocal. Relative intensity was calculated for each channel using ImageJ. Images were subjected to *maximum projection* and background subtraction using the *rolling ball* function, quantified using ROIs for single cells, and normalized to control. Data are presented with the SEM from three independent experiments, with statistical significance determined by a t-test (***p < 0.001, ****p < 0.0001). **B.** Flow cytometry analysis (FACS) of the Clover and mScarletI fluorescence response in duplex endo-tagged GABARAPs to autophagy-inducing conditions. The average mean fluorescence intensity was measured in fluorescence units and presented with the SEM from three independent experiments. Statistical significance was determined by a t-test, with **p < 0.01, ***p < 0.001, ****p < 0.0001, ns – insignificant.

Our endo-tagged duplex reports may thus faithfully phenocopy their single endo-tagged counterparts, combined with the added value of simultaneous characterization of distinct localization patterns of specific isoforms.

### Isolation of distinct autophagic vesicles from endo-tagged GABARAPs cells

Recent studies have highlighted distinct roles for individual GABARAPs in autophagy and other cellular processes [11, 19]. These insights align with our observations of predominantly unique localizations for GABARAPs. Consequently, we hypothesized that different GABARAPs are associated with distinct autophagic vesicles. To investigate this, we employed the 3xFlag epitope to immunoprecipitate and isolate GABARAPs-positive membranes obtained from cells under basal conditions (*Figure 4A*).

**Figure 4.**
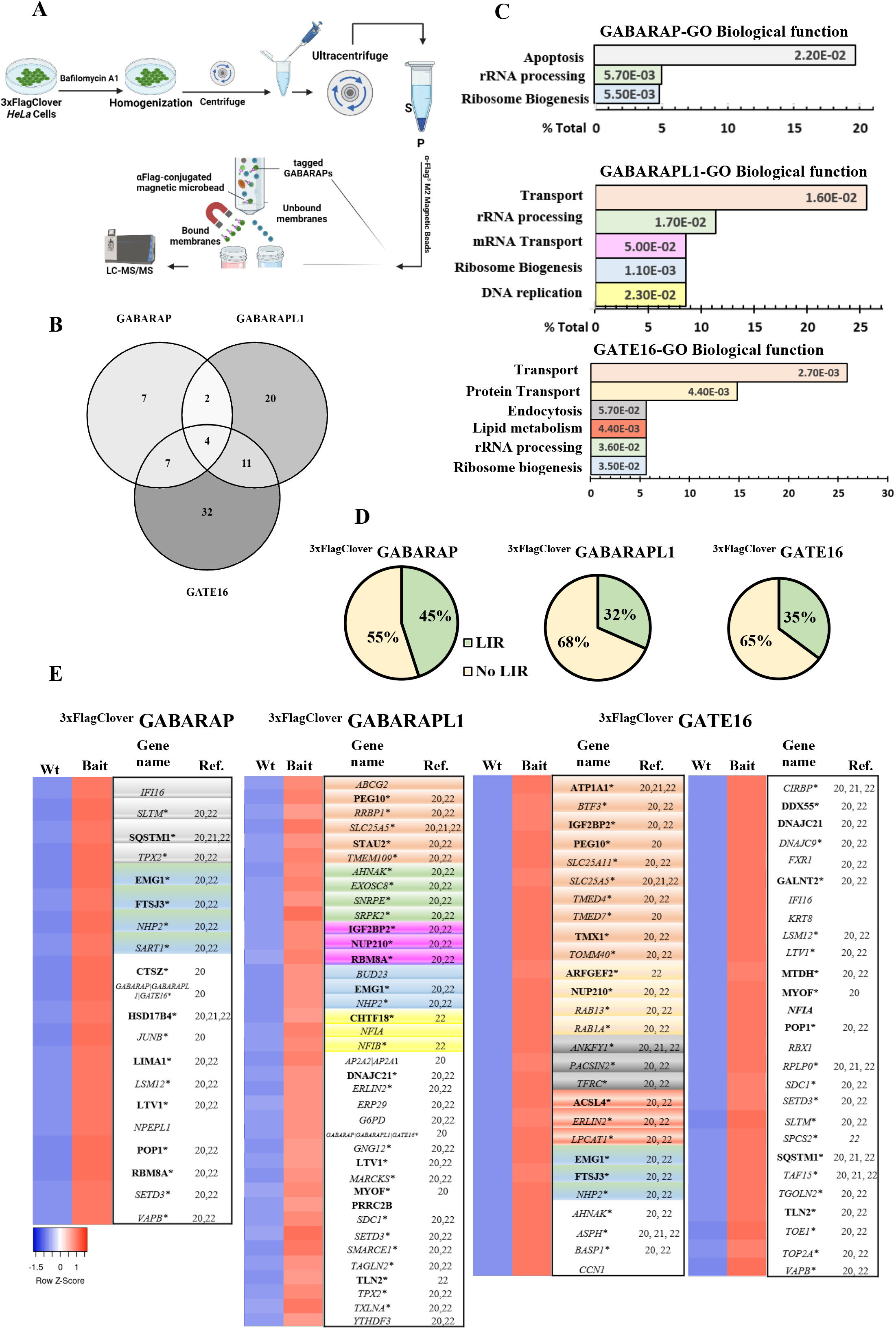
Isolation of distinct autophagic vesicles from endo-tagged GABARAPs cells. **A.** Schematic representation of the immunoprecipitation process for isolating single-color endo-tagged GABARAP-positive membranes using Anti-FLAG M2 Magnetic Beads. Parental *HeLa* cell lines and endo tagged GABARAPs reporter cells were treated with 0.1 μM Bafilomycin A1 (Baf.A1) for 2 hours, homogenized by sonication (3 pulses of 3 seconds each at 60% amplitude), and centrifuged to remove cell debris, followed by ultracentrifugation. The membrane fraction was then incubated with Anti-FLAG M2 Magnetic Beads for 2 hours for immunoprecipitation under non-reducing conditions. The GABARAP-positive membranes underwent mass spectrometry analysis. Figure created with BioRender.com **B.** Overlap of proteins found between and among GABARAP subfamilies as determined by LC-MS/MS. **C.** DAVID gene ontology (GO) analysis of proteins enriched more than 2-fold with a p-value < 0.05.**D.** LIR domain analysis of significant hits from the LC-MS/MS, classified by the iLIR online server (https://ilir.warwick.ac.uk/) using PSSM score prediction. Proteins with a PSSM score >16 were classified as LIR-containing. **E.** Subsets of enriched proteins in LC-MS/MS analysis of the GABARAP subfamily (Supplementry Table 2 for the complete data list), colored according to the GO function analysis clusters in presented in **Figure 4C**. Proteins lacking clustering are shown in white. Proteins containing LIR domains are highlighted in bold, while those without LIR domains are presented in Italics. Asterisks denote proteins found in other autophagy-related proteomics studies [20–22].

Mass spectrometry-based proteomic analysis of the eluates identified in duplicate experiments 112 proteins whose abundances in GABARAP immunoprecipitates were higher by 2-fold than the untagged control (*Figure 4*). Notably, 82% of these proteins have been previously reported in other autophagy-related studies [20–22] (*Figure 4E*). The presence of LC3 interaction region (LIR motifs) was detected on approximately one-third of the proteins found in the isolated GABARAPs specific vesicles (45% for GABARAP, 32% for GABARAPL1, and 35% for GATE16) (*Figure 4D*), confirming the potency of our isolation method in enrichment of autophagic membranes and the successful application of the affinity tag. Functional annotation using the DAVID platform [23] revealed their involvement in vesicular trafficking, protein transport, and mRNA transport, as well as in RNA processing, lipid metabolism, ribosome biogenesis, and apoptosis. These, too, are consistent with previous reports on autophagosomal general content [21]. Intriguingly, the pathways varied between different GABARAPs in type and prominence (*Figure 4C*). Only four of the identified proteins found were shared across all three GABARAP homologs, suggesting distinct content for each autophagosome labeled by the different GABARAP, and possibly highlighting functional differences among GABARAP subfamily proteins (*Figure 4B*).

## Discussion

Atg8 proteins are central to autophagy [1]. In yeast, a single Atg8 protein mediates selective and nonselective autophagy, while higher eukaryotes express up to seven family members in mammals and nine in plants [24, 25]. Research on the exact function of individual Atg8 proteins in this process is currently limited, mainly due to a lack of cellular tools. The Atg8 family is dissected into two subfamilies, LC3 and GABARAP, both needed for autophagy [26]. GABARAP subfamily members have been described as pivotal in autophagy and other cellular processes [10]. Here, we describe a new cellular platform for studying the functions of individual GABARAPs in autophagy and possibly other cellular pathways.

Numerous assays are currently available to study autophagy, greatly enhancing our understanding of its mechanisms. LC3B, an ATG8 member, is a widely used marker for measuring autophagic flux and activity [14]. Much of our present understanding of human Atg8 orthologues stems from studies on LC3B, often applying its findings to the entire family. Growing evidence indicates functional variations among the GABARAPs and LC3s subfamilies, pointing out GABARAPs unique roles in bulk and selective autophagy [10, 26, 27]. However, the high sequence and structural similarity within and across the LC3 and GABARAP subfamilies [20] requires the development of particular and sensitive readout systems for their precise and non-redundant functions. Current autophagy studies primarily rely on ectopically expressed Atg8s, which may falsely modulate the autophagic flux [25] or form an autophagy-independent aggregates [28]. Furthermore, autophagy high-throughput studies using overexpression systems are prone to low sensitivity due to the unnatural abundance of overexpressed ATG8 proteins, frequently requiring additional preparatory steps analysis [14, 29]. CRISPR-Cas9 endogenous tagging of mammalian ATG8 proteins has been thus far limited to biochemical and fixed-sample studies, which are prone to biases and only provide a snapshot of a dynamic process [12, 20]. Endogenous tagging of LC3 and GABARAP with fluorescent proteins has uncovered new autophagy modulators [30] and interactions [31], demonstrating the potential of such systems. However, the low abundance of ATG8s limits fluorescence necessitates complementation with transient expression of ATG8s or immunostaining [29], and typically focuses on a single ATG8 member [30]. Our approach, employing Clover [15] and mScarletI [16] fluorescent proteins, far brighter than standard GFP or RFP, enables simultaneous visualization and high-throughput analysis of GABARAP homologs. Furthermore, including 3xFlag epitope-tagged proteins enables comprehensive and complementary biochemical and proteomic analyses. Nevertheless, our assay system suffers from certain limitations, notably its specificity to a single cell type, unlike transient expression methods adaptable to various cells. In addition, flux estimation in this study relies on chemical inhibition of lysosomal degradation with Bafilomycin-A1 instead of direct measurements of autophagic clearance of GABARAP proteins in pulse and chase experiments [29, 30, 32].

In summary, this study introduces a novel platform for investigating the GABARAP subfamily’s autophagy roles, using cell lines that facilitate live visualization and high-throughput analysis with brighter fluorescent proteins, complemented by comprehensive biochemical and proteomic analyses. Notably, in real-time, duplex cell lines reveal distinct vesicle localizations for GABARAP-occupied vesicles. Coupled with proteomic data, these observations indicate functional variations among GABARAP homologs, prompting a reconsideration of autophagic research’s focus toward a homolog-dependent approach and providing a platform for such studies within the GABARAP subfamily.

## Acknowledgments

Z.E. is the incumbent of the Harold Korda Chair of Biology. We are grateful for funding from the Israel Science Foundation (Grant 215/19), Joint NRF - ISF Research Fund (Grant 3221/19), Joint NSFC-ISF Research Fund (Grant 3345/20), and the Yeda-Sela Center for Basic Research. We thank Renana Shvimmer, Shahar Lyubinsky, Shahar Vanunu and Michal Levi for their technical assistance. We also thank Drs. Yifat Merbl, Yaron Antebi and Oren Shatz for their insightful comments and suggestions. Thanks to Dr. Yossef Addadi, Tatyana Smirnova, and Inna Goliand from the Advanced Optical Imaging Unit, de Picciotto-Lesser Cell Observatory unit at the Moross Integrated Cancer Center Life Science Core Facilities, Weizmann Institute of Science for their scientific support and discussions.

## Methods

### Cell cultures and treatments

*HeLa* parental cells (strain JW; obtained from the Weizmann Institute Cell-Line Core facilities) were cultured in alpha minimum essential media (αMEM; Biological Industries, 01-042-1A) supplemented with 10% fetal calf serum (FBS; Invitrogen, 10270106), 100 IU/mL penicillin, and 100 μg/mL streptomycin at 37°C in a 5% CO2 environment. For the induction of autophagy, cells were washed twice with phosphate-buffered saline (PBS; Biological Industries, 02-023-1A) and then incubated in starvation medium Earle’s Balanced Salt Solution (EBSS; Biological Industries, 02-011-1A) for 4 hours. Lysosomal degradation was inhibited using 100 nM Bafilomycin A1 (LC Laboratories, B-1080) for 4 hours. To ensure quality, all cell lines were routinely checked for mycoplasma contamination on a monthly basis.

### Plasmids

For donor plasmid construction, Left and Right Homology Arms (1000 bp each) were PCR-amplified with the genomic DNA of *HeLa* parental cells. Clover and mScarletI sequences were obtained from Addgene plasmids 40259 and 85044, kindly provided by Yosef Shaul and Eitan Reuveny. To circumvent potential steric hindrance from the tags, a Glycine-Serine (GGGS)3 linker was introduced between the fluorescent protein and GABARAPs within the integration cassette (*Figure 1A*), designed using SynLinker [33], an online tool for predicting minimal energy scores (kJ/mol) and assessing steric clashes. All fragments, including the integration cassette flanked by Left (HA-L) and Right (HA-R) Homology arms (*Figure 1A*), were assembled into the pBlueScript KS (NovoPro V011756) plasmid backbone, also a gift from Yosef Shaul, using GeneArt™ Gibson Assembly HiFi Master Mix (Thermo Fisher Scientific, A46628). The 3XFlag sequence, synthesized as a gBlock by IDT was integrated into the donor plasmid through restriction-free cloning. A synonymous mutation was introduced at the PAM site in HA-R to prevent Cas9 re-cleavage post-integration (*Supplementary Table 1*.)

Single-guide RNAs (sgRNAs) for endogenous tagging and ATG14 knockout were designed using Benchling online platform [https://benchling.com] and inserted via BsaI sites into the pU6-(BsaI)_CBh-UN-Cas9 plasmid (Addgene 135011) [34] for endogenous tagging and BbsI sites forpSpCas9(BB)-2A-Puro (PX459) V2.0 plasmid (Addgene 62988), for ATG14 knockouts. The LAMP1-RFP plasmid (Addgene plasmid 1817), kindly provided by Ori Avinoam, and a custom LAMP1-iRFP670 plasmid, developed by substituting RFP with the iRFP670 sequence from CMV-H2A-iRFP670 (courtesy of Ravid Straussman), were utilized. Details of the primers and sgRNAs are listed in *Supplementary Table 1*.

### CRISPR–Cas9 genome editing

For endogenous tagging, we employed a chimeric construct of Cas9, in which SpCas9 was fused at its N-terminus to the intrinsically disordered viral protein (IDP), enhancing the recruitment of the MRN complex (Mre11/Rad50/NbsI), crucial for homology-directed repair, thereby improving genome integration events. [34]. *HeLa* cells were transfected with the designatedpU6-(BsaI)_CBh-UN-Cas9 plasmid expressing sgRNA targeting a specific GABARAP subfamily member, along with the corresponding donor plasmid. Five days post-transfection, cells expressing a fluorescent tag were selected and sorted into single-cell populations by FACS to 96 well plates. The successful integration of the tag was confirmed by PCR, sequencing, and immunoblotting.

Knockout of ATG14 in single-tagged GABARAPs reporter cells was achieved by transfecting the desired cell line with pSpCas9(BB)-2A-Puro (PX459) V2.0 plasmid (Addgene 62988) expressing ATG14-targeting sgRNA. Seventy-two hours post-transfection, cells were washed with PBS, trypsinized, and replated in culture media containing 3 mg/ml puromycin. After 48 hours, surviving cells were sorted into single-cell populations by limiting dilution to 1 cell per well per 96-well plates. Knockout clones were identified via sequencing. The absence of ATG14 transcripts was verified by Reverse Transcription PCR (RT-PCR) on cDNA synthesized from total RNA extracted from the cell samples using Terra™ PCR Direct Polymerase Mix (TAKARA, 639270). Total RNA was isolated using the NucleoSpin RNA II kit (Macherey-Nagel), and 1 μg of RNA was reverse-transcribed using M-MLV reverse transcriptase (Promega) with random hexamer primers (Amersham).

### Flow cytometry detection of gene-edited cells

Fluorescence-activated cell sorting was used to generate endogenously tagged GABARAP cell lines. Cells were transfected with the appropriate pU6-(BsaI)_CBh-UN-Cas9 and donor plasmids. Five days post-transfection, cells were treated for 2 hours with Baf.A1, then washed with PBS and detached by trypsinization. The cell pellet was collected by centrifugation (3 minutes at 1000 g), resuspended in ice-cold PBS with 2% (v/v) FBS to 1 × 10^6 cells, and kept on ice before sorting. Cells endogenously expressing fluorescent tags were selected and sorted into 96 wells. *HeLa* wt cells were used as a negative control to distinguish between nonspecific and specific FP/Clover/mScarletI expression levels. Cells were analyzed and sorted based on fluorescence signal relative to the unstained parental control using BD FACSAria II (BD Biosciences, Franklin Lakes, NJ).

Flow cytometry analysis of endogenously tagged *HeLa* cells was done as described previously [18]. Cells were seeded at 80% confluence in 96-well plates and starved using EBSS media with and without Baf.A1, for 4 hours prior to analysis. Following starvation, cells were washed with PBS, detached by trypsinization, the pellet was washed with PBS by centrifugation (3 minutes at 1000 g), and then resuspended in ice-cold PBS maintained on ice before analysis.

Fluorescent proteins were excited using 488nm laser line 488 nm (mClover3) and 532 nm laser line (mScarletI), and emissions were detected at maxima/using bandpass values filters of 525/50 nm and 575/25 nm, respectively. A total of 1 × 105 events were analyzed per sample, with unmodified *HeLa* cells serving as a control. Autofluorescence baselines (<1 × 10^3 fluorescent units) for relevant parameters were established with non-modified *HeLa* cells. Data analysis was performed using FlowJo software v10.7.2 (FlowJo LLC, Ashland, OR).

### Knockdown of GABARAP family proteins

Endogenously tagged GABARAP subfamily reporter cells were transfected with siRNA SMARTpool sequences at a final concentration of 50 nM using DharmaFect 1 (Dharmacon) according to the manufacturer’s instructions. The siRNA oligos (Dharmacon), comprising four RNA duplexes, were targeted to GABARAP (M-012368-01), GABARAPL1 (M-014715-01), GATE-16 (M-006853-02), and included a non-targeting control (D-001206-14).

### Immunoblotting

Total cellular protein extracts were prepared using RIPA buffer (0.1 M NaCl [Bio-Lab Ltd, 21955], 5 mM EDTA [J.T. Baker, 8993], 0.1 M sodium phosphate [Sigma, 342483], pH 7.5, 1% Triton X-100 [Sigma, X100], 0.5% sodium deoxycholate [Sigma, D6750], 0.1% sodium dodecyl sulfate [Sigma, L4509]), supplemented with a protease inhibitor cocktail (PIC; Merck, 539134). Extracts were centrifuged at 16,000 x g for 15 minutes at 4°C, and protein concentrations were determined using Bio-Rad Protein Assay Dye Reagent Concentrate (Bio-Rad, 500-0006). Proteins were then separated by SDS-PAGE using an 8-16% polyacrylamide gradient gel (Invitrogen) and transferred onto a nitrocellulose membrane (Bio-Rad, 1704159). For immunoblotting, the samples were transferred from the SDS-PAGE gel to Trans-Blot Turbo Midi 0.2 µm Nitrocellulose (Bio-Rad, 1704159) with Trans-Blot Turbo Transfer System (Bio-Rad). After incubation with the relevant antibody, the signals from incubation with Enhanced ChemiLuminescence (ECL) detection system (Biological Industries, 20-500-120 were detected with AmerchamTM Imager 680 (Cytivia). Band intensities were measured with a Gel Analyzer in the open-source image processing software ImageJ (version 1.54)

### Live imaging and Image analysis of endogenously tagged GABARAPs

Live *HeLa* cells were seeded onto a 24-well glass-bottom black plate (Cellvis, P24-1.5H-N) at least 48h before imaging. For autophagy induction, cells were starved 4h before imaging. Cells were incubated with 100 nM Bafilomycin A1 for 4 hr before imaging to inhibit lysosomal degradation. The incubation chamber held at 37°C and 5% CO2 during imaging.

Live imaging was conducted using a Dragonfly 505 spinning disk confocal microscope (Andor Technology PLC) spinning disk confocal microscope with 40 um pinhole size disk equipped with an Andor Zyla-4.2P sCMOS camera and Leica HC PL APO 63x/1.30 GLYC CORR CS2 objective (11506353). The incubation chamber held at 37°C and 5% CO2 during imaging. The fluorescent proteins were detected using the following excitation and emission filter combinations: Clover (Ex-488, Em-525/50), mScarlet-I/Lamp1-RFP (Ex-561, Em-600/50), and Lamp1-iRFP670 (Ex-640, Em-700/75). The microscope was operated using Fusion 2.3 software, and image analysis was performed with ImageJ (version 1.54). Zstacks with step size in the range 0.166 – 0.288 um with 200-400 ms exposure.

For Airyscan super-resolution microscopy, cells were imaged in FluoroBrite DMEM (Thermo Fisher Scientific) with 10% (v/v) FBS (Corning Corp.). Airyscan imaging was performed using a Zeiss 880 (Carl Zeiss AG, Oberkochen, Germany) outfitted with an Airyscan module and incubation chamber held at 37°C and 5% CO2. Data were collected using a 63 × 1.4 NA objective and immersion oil optimized for 37°C (Carl Zeiss AG). Colors were collected sequentially to minimize crosstalk at a pixel resolution of 0.04 μm, 2-4 sec/Frame. Airyscan processing performed using the Airyscan module in the commercial ZEN black software package (Carl Zeiss AG).

### Immunofluorescence

For p62 colocalization analysis, cells cultured on 96-well glass bottom plates (Cellvis, P96-1.5H-N) and treated as indicated were fixed and permeabilized using 100% methanol (Bio-Lab Ltd, 136805) for 10 minutes at −20°C. Following fixation, cells were blocked with 10% FCS in PBS for 30 minutes at room temperature. Then incubated with the primary antibody diluted in 2% FCS in PBS for 1 hour at room temperature followed by a secondary antibody, supplemented with 1 μg/ml Hoechst 33342 (Invitrogen, H1399) in 2% FCS in PBS, for 30 minutes at room temperature. Imaging was performed using a Dragonfly 505 (Andor Technology PLC) spinning disk confocal microscope with 40 um pinhole size disk equipped with an Andor Zyla-4.2P sCMOS camera and Leica HC PL APO 63x/1.30 GLYC CORR CS2 objective (11506353), resulting pixel size 0.096 um and step size in the range 0.166 – 0.288 um. The fluorescent proteins and antibodies were detected using the following excitation and emission filter combinations: Hoechst 33342 (Ex-405, Em-450/50), mClover3 (Ex-488, Em-525/50), mScarlet-I (Ex-561, Em-600/50), and Cy5 (Ex-640, Em-700/75). The microscope was operated using Fusion 2.3 software, and image analysis was performed with ImageJ (version 1.54).

### Mass-spectrometry analysis and immunoprecipitation of GABARAPs positive membranes

Single reporter cells and parental controls were grown to confluence and treated with Bafilomycin A1 (Baf.A1) for 2 hours to enrich the autophagosomal fraction. Subsequently, cells were homogenized via sonication, employing three cycles of 3-second pulses at 60% amplitude. Cell debris was removed by centrifugation at 10,000 x g for 10 minutes at 4°C. The resulting supernatants were then ultra-centrifuged at 90,000 rpm (using rotor TLA120) for 45 minutes at 4°C. The resultant pellet fraction, containing autophagosomes, was resuspended in homogenization buffer (10 mM Tris pH 7.5, 0.3 M Sucrose, 50 mM KCl) supplemented with two mM PMSF, two mM NEM, and a protease inhibitor cocktail. Following the manufacturer’s instructions, immunoprecipitation was performed with Flag magnetic beads (Sigma; M8823).

### Sample preparation

Proteins were eluted with 5% SDS in 50 mM Tris-HCl. Proteins were reduced with five mM dithiothreitol at 60°C for 45 minutes and alkylated with ten mM iodoacetamide in the dark for 45 minutes. Each sample was loaded onto S-Trap microcolumns (Protifi, USA) according to the manufacturer’s instructions. In brief, after loading, samples were washed with 90:10% methanol/50 mM ammonium bicarbonate. Samples were then digested with 250ng trypsin for 1.5 h at 47 °C. The digested peptides were eluted using 50 mM ammonium bicarbonate. An additional round of trypsin was added to this fraction and incubated overnight at 37 °C. Two more elutions were made using 0.2% formic acid and 0.2% formic acid in 50% acetonitrile. The three elutions were pooled together and vacuum-centrifuged to dry. Samples were kept at −80 °C until analysis.

### Liquid chromatography

ULC/MS grade solvents were used for all chromatographic steps. Each sample was loaded using split-less nano-Ultra Performance Liquid Chromatography (Acquity M Class; Waters, Milford, MA, USA). The mobile phase was A) H_2_O + 0.1% formic acid and B) acetonitrile + 0.1% formic acid. The samples were desalted online using a reversed-phase Symmetry C18 trapping column (180 µm internal diameter, 20 mm length, 5 µm particle size; Waters). The peptides were then separated using a T3 HSS nano-column (75 µm internal diameter, 250 mm length, 1.8 µm particle size; Waters) at 0.35 µL/min. Peptides were eluted from the column into the mass spectrometer using the following gradient: 4% to 30%B in 55 min, 30% to 90%B in 5 min, maintained at 90% for 5 min, and then back to initial conditions.

### Mass Spectrometry

The nanoUPLC was coupled online through a nanoESI emitter (10 μm tip; Fossil, Spain) to a quadrupole orbitrap mass spectrometer (Q Exactive HF Thermo Scientific) using a FlexIon nanospray apparatus (Thermo Scientific). Data was acquired in Data Dependent Acquisition mode, using a Top10 method. MS1 resolution was set to 120,000. Mass range 375-1,650 Th. Maximum injection time of 60msec, automatic gain control (AGC) was set to 1e6. MS/MS resolution was set to 15,000. Injection time of 60msec, AGC 1e5, NSE of 27, and dynamic exclusion of 20sec.

### Data processing

The raw data processing was performed using MetaMorpheus version 0.0.320, available at https://github.com/smith-chem-wisc/MetaMorpheus. The following search settings were used: protease = trypsin; maximum missed cleavages = 2; minimum peptide length = 7; maximum peptide length = unspecified; initiator methionine behavior = Variable; fixed modifications = Carbamidomethyl on C, Carbamidomethyl on U; variable modifications = Oxidation on M; max mods per peptide = 2; max modification isoforms = 1024; precursor mass tolerance = ±5 PPM; product mass tolerance = ±20 PPM. The combined search database contained 20537 non-decoy protein entries, including 296 contaminant sequences. The database was downloaded from UniProtKB. The resulting output was loaded to Perseus version 1.6.2.3. Data was Log2 transformed. Proteins were filtered for replication in at least 2 of 3 replicates. Missing data was imputed from a low, random distribution with the default values. For functional annotations, the platform DAVID (https://david.ncifcrf.gov/) was used. The mass spectrometry proteomics data have been deposited to the ProteomeXchange Consortium via the PRIDE [35] partner repository with the dataset identifier PXD049031.

### Statistical analysis

Where appropriate, statistical significance between data sets was analyzed by *t*-tests using GraphPad Prism (version 10.1.1, GraphPad Software, San Diego, CA, USA). ns. - non-significant; *p<0.05, **p<0.01, ***p<0.001, ****p<0.0001.

## Supplementary figure legends

**Figure 1. Figure Supplement 1. Generation of single and duplex GABARAPs reporter cell lines. A.** GABARAPs single color reporter cells were transfected with nontargeting (control) siRNA (si*NT*) or GABARAP or GATE16 or GABARAPL1 siRNA using DharmaFECT1 transfection reagent for 72 h. Total protein extracts were analyzed by western blotting for Flag, GFP, and GPDH (loading control). **B.** GABARAPs duplex reporter cells were transfected with nontargeting (control) siRNA (si*NT*), or with siGABARAP, siGATE16, or siGABARAPL1, using DharmaFECT1 transfection reagent for 72 h. Total protein extracts were analyzed by western blotting for Flag, GFP, RFP, and GPDH (loading control).

**Figure 2. Figure Supplement 1. Analyzing autophagy by single-tagged GABARAP reporter cells. A.** Quantification of SQSTM1 (p62) and ^3xFlag-Clover^GABARAPs flux based on p62 and Flag antibody respectively, as depicted in Figure 2C. Levels of SQSTM1 (p62) and Flag with the SEM of three independent experiments, *p < 0.05, **p < 0.01, were determined by t-test. **B.** Reverse PCR (RT PCR) for ATG14 gene. Cells were grown to confluence, followed by total RNA extraction and reverse transcription. cDNA was used as a template for PCR, using primers targeting the ATG14 transcribed region. C. Single endo tagged GABARAPs response to autophagy-inducing conditions with and without ATG14, indicated by western blot analysis. Cells were grown to confluence in a complete and starvation (EBSS) medium and treated (where indicated) for the last 4 h with 0.1 µM Bafilomycin A1. Total protein extracts were probed for Flag, GFP, SQSTM1 (p62), and GPDH (loading control).

**Figure 2. Figure Supplement 2 Analyzing autophagy by single-tagged GABARAP reporter cells. A.** Representative images of single-color endo-tagged GABARAPs and the lysosomal marker LAMP1 by Airyscan super-resolution microscopy. All three reporter cell lines were transfected with the LAMP1-RFP plasmid using JetPrime reagent for 48 hours. Subsequently, cells were incubated for 4 hours in a control medium (DMSO) or starvation medium (EBSS) and treated with 0.1 μM Bafilomycin A1 were indicated for 4 hours. Scale bar: 10µm, in Zoom figures 1µm. For analysis, the visualization was performed using scanning confocal microscopy. Colocalization was quantified by Pearson correlation coefficient for LAMP1-RFP and GABARAPs, calculated using ROIs for single cells by *Coloc2* module with 10 *Costes* iterations in ImageJ, and data are presented with the SEM from three independent experiments. Statistical significance was determined by a t-test, with ****p < 0.0001. **B.** Single color endo-tagged GABARAPs cells were incubated in complete medium or EBSS in the presence of 0.1 µM Bafilomycin A1 where indicated, for 4 h. Then cells were fixed in absolute methanol and immunostained with SQSTM1 (p62) antibody. The visualization was performed using spinning disk confocal microscopy. Scale bar 10µm. Colocalization was quantified using ROIs for single cells, by Pearson correlation coefficient for SQSTM1 (p62) and GABARAPs, using *Coloc2* module with 10 costers iterations at ImageJ and presented with the SEM of three independent experiments, ****p < 0.0001 determined by t-test.

**Figure 3. Figure Supplement 1. Measuring autophagy with the duplex GABARAPs reporter system. A.** Double-tagged GABARAPs response to autophagy-inducing conditions indicated by western blot analysis. Cells were grown to confluence in a complete and starvation (EBSS) medium and treated (where indicated) for the last 4 h with 0.1 µM Bafilomycin A1. Total protein extracts were probed for Flag, SQSTM1 (p62), GFP, RFP and GPDH (loading control). Levels of Flag and SQSTM1 (p62) with SEM of three independent experiments, *p < 0.05, ***p < 0.001 were determined by t-test. **B.** Representative Airyscan super-resolution images for duplex GABARAPs reporter cell lines, scale bar 10µm, in Zoom figures 1µM. For image analysis, visualization was done using spinning disc confocal. Clover and mScarletI overlap quantified using Manders correlation coefficient. Image analysis employed the *Coloc2* module with 10 Costes iterations in ImageJ and background subtraction using the *rolling ball* function, quantified using ROIs for single cells.

**Figure 3. Figure Supplement 2. Measuring autophagy with the duplex GABARAPs reporter system.** Duplex GABARAPs reporter cell lines were incubated in complete medium or EBSS in the presence of 0.1 µM Bafilomycin A1 where indicated, for 4 h. Then cells were fixed in absolute methanol and immunostained with SQSTM1 (p62) antibody. The visualization was performed using spinning disk confocal microscopy. Scale bar 10µm. Colocalization was quantified using the Pearson correlation coefficient for p62 and GABARAPs, employing the *Coloc2* module with 10 Costes iterations in ImageJ, using ROIs for single cell. Data are presented with the SEM from three independent experiments. Statistical significance was determined by a t-test, with ***p < 0.001, ****p < 0.0001.

**Figure 3. Figure Supplement 3. Measuring autophagy with the duplex GABARAPs reporter system.** Duplex GABARAPs reporter cells exogenously expressing Lamp1-iRPF670 were incubated in complete medium or EBSS in the presence of 0.1 µM Bafilomycin A1 where indicated, for 4 h. The visualization was performed using spinning disk confocal microscopy. Scale bar 10µm. Colocalization was quantified using the Pearson correlation coefficient for iRFP670 and Clover/mScarlet puncta, employing the *Coloc2* module with 10 Costes iterations in ImageJ, using ROIs for single cells. Data are presented with the SEM from three independent experiments. Statistical significance was determined by a t-test, with ****p < 0.0001.

**Supplementry table 1.**
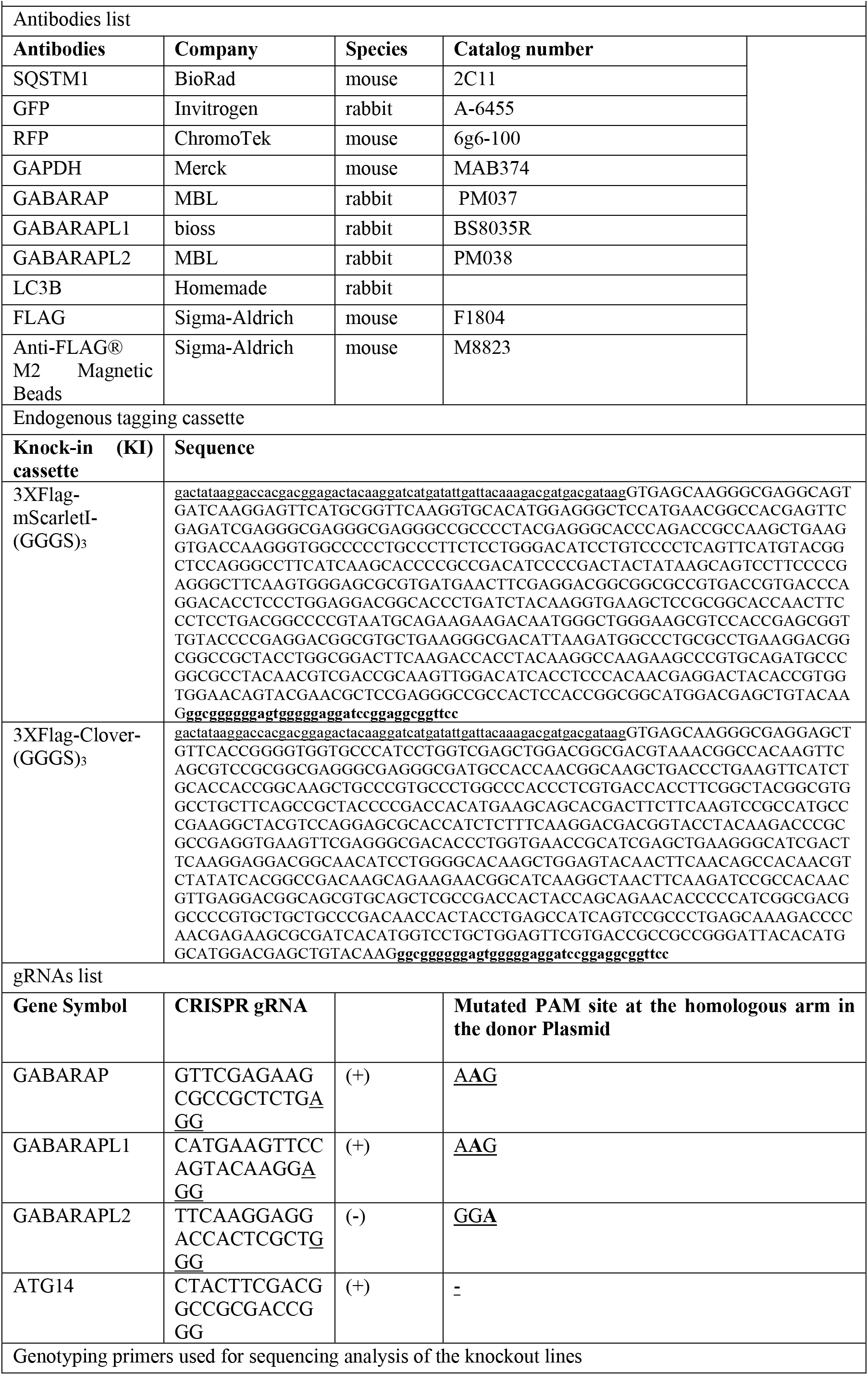
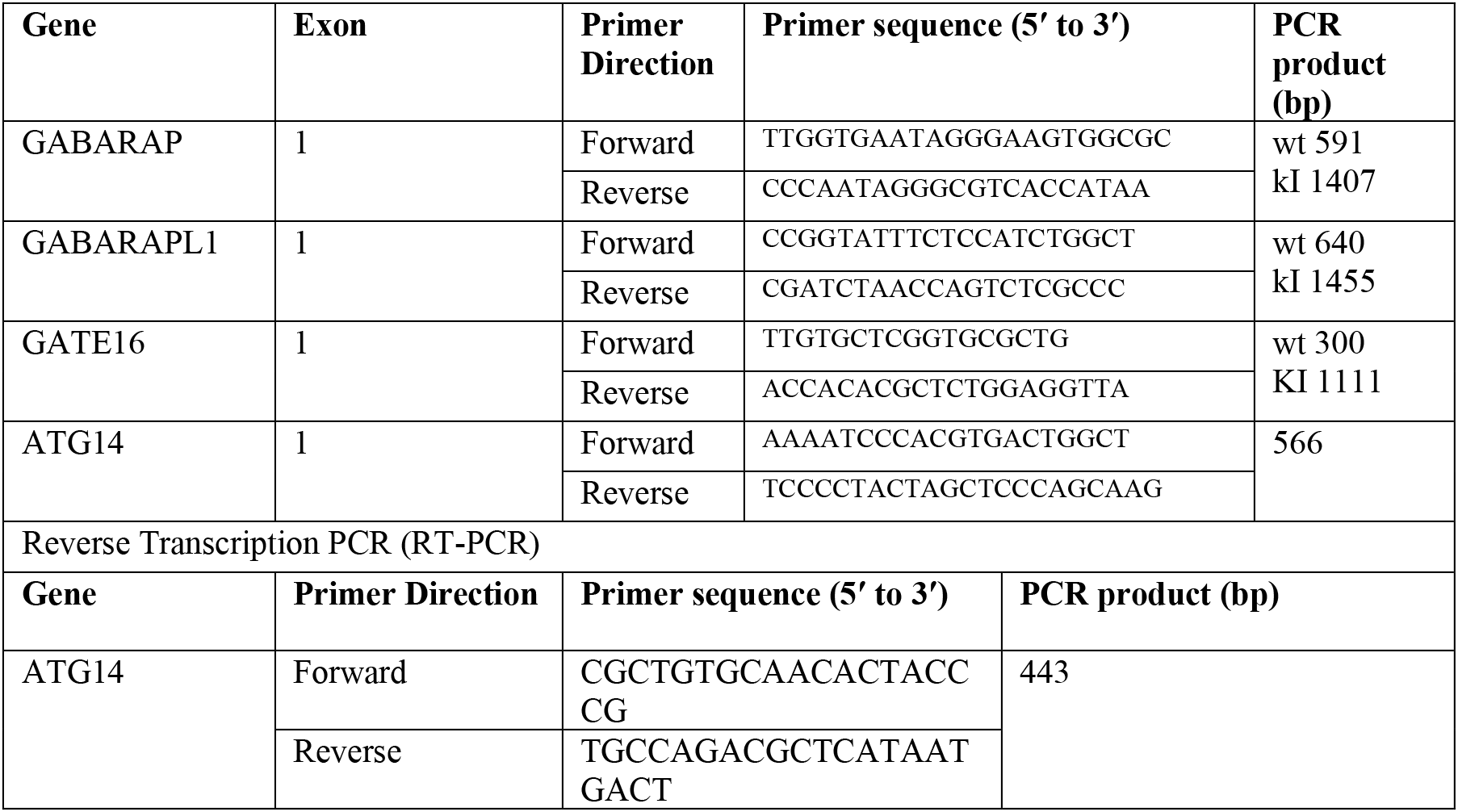

