## Supplementary figures and images for "A new cellular platform for studying autophagy"

### Figure 1 Supplement figure 1

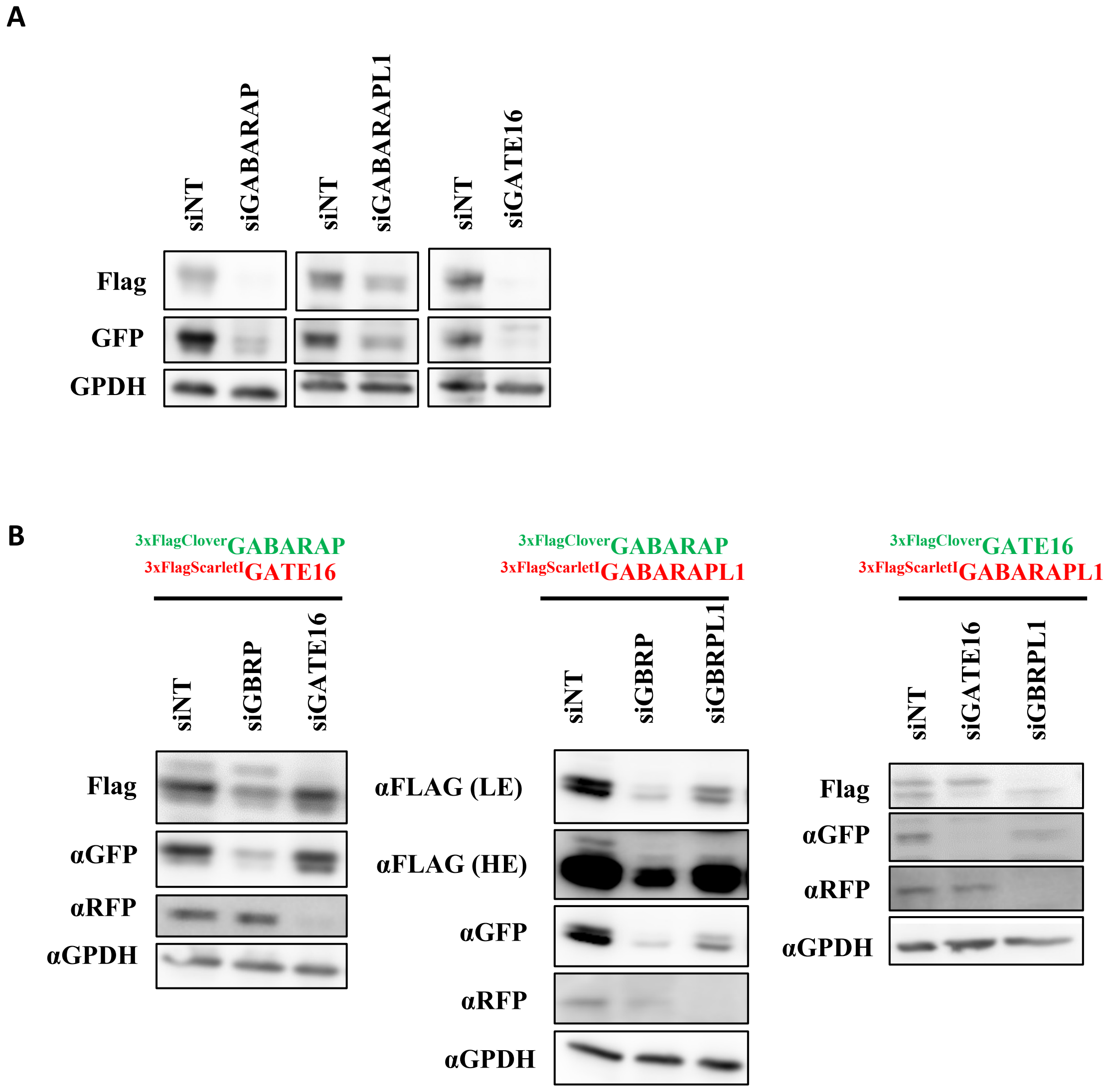


**Figure 1. Figure Supplement 1.**

### Figure 2 Supplement figure 1

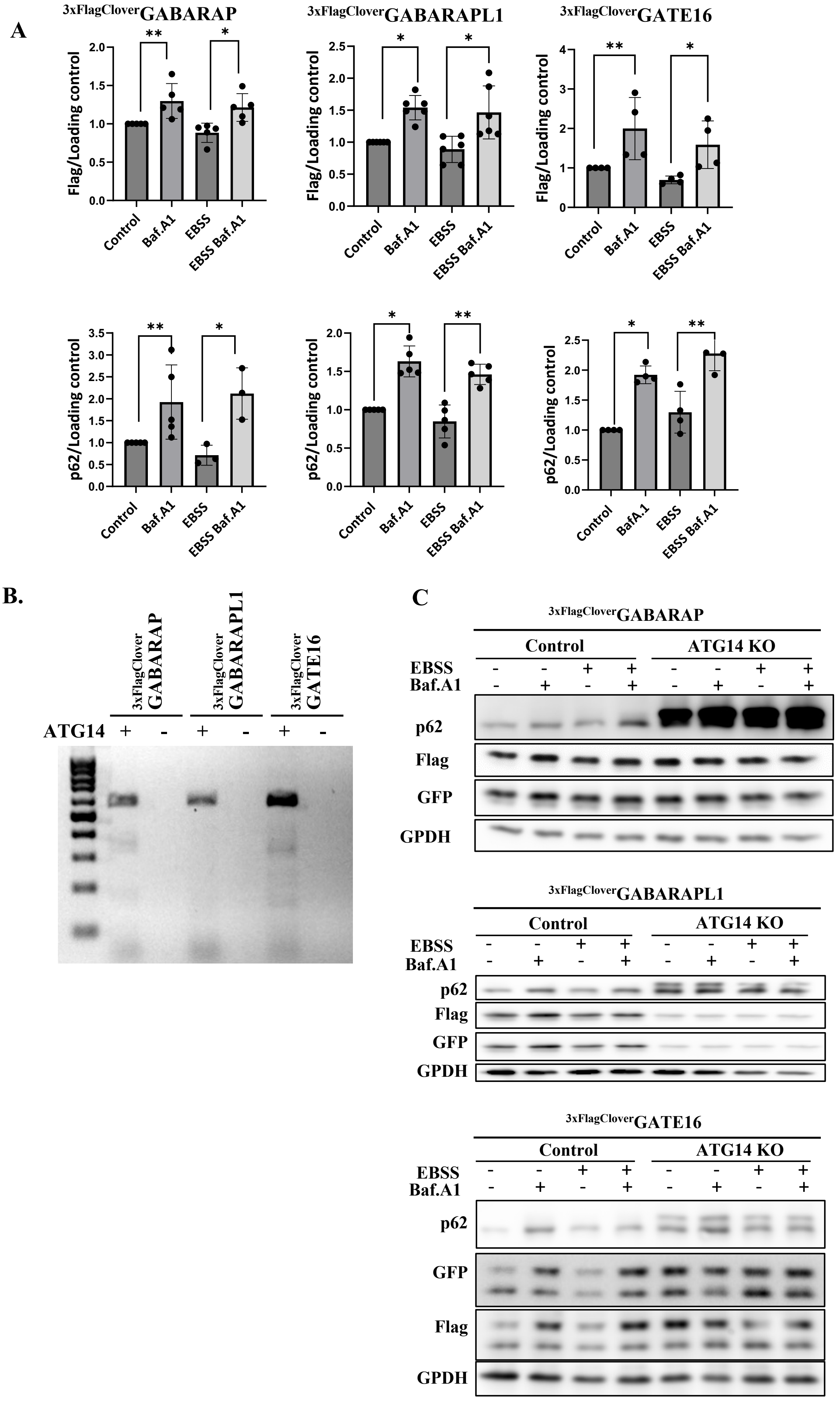


Figure 2. Figure Supplement 1.

### Figure 2 Supplement figure 2

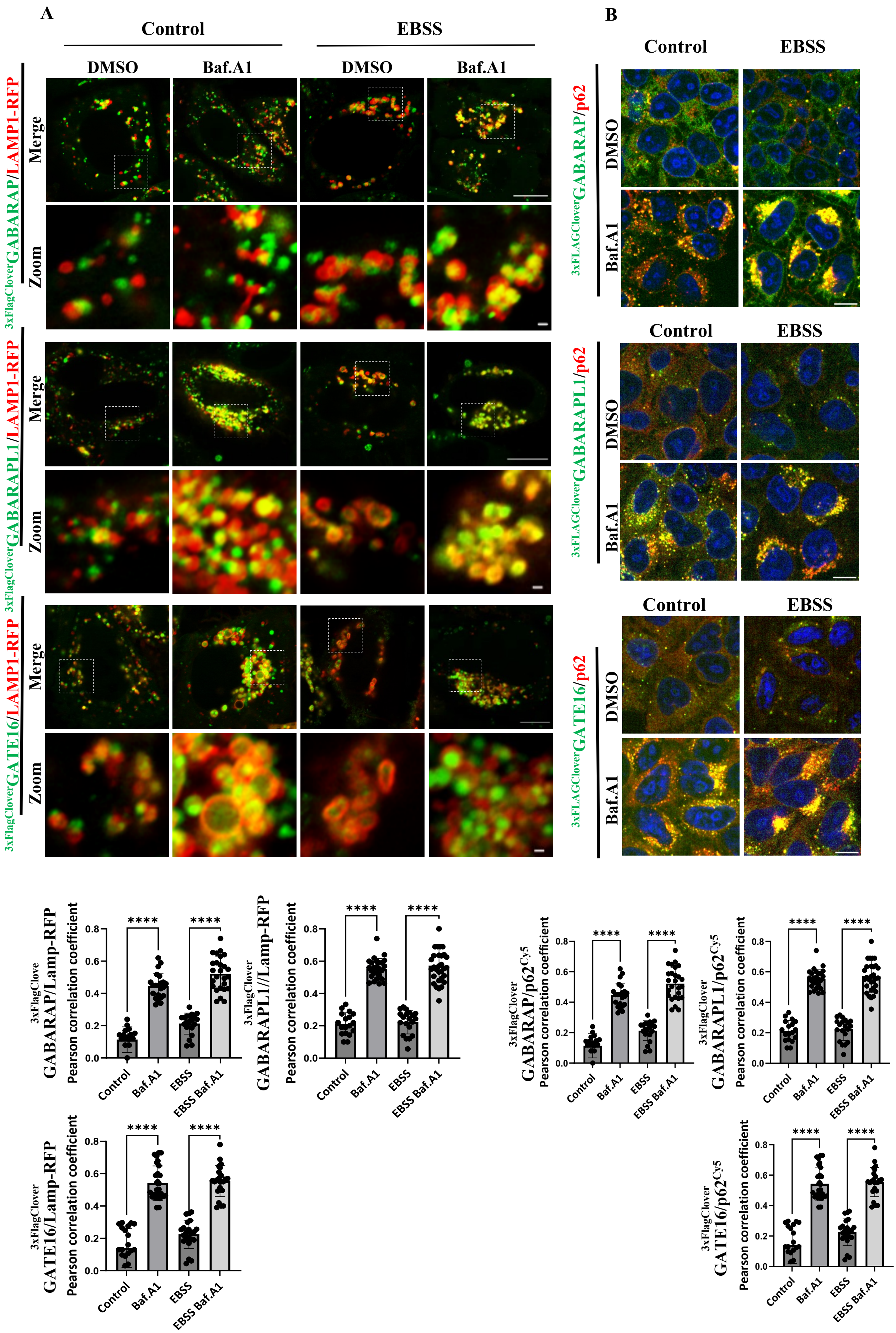


**Figure 2. Figure Supplement 2.**

### Figure 3 Supplement figure 1

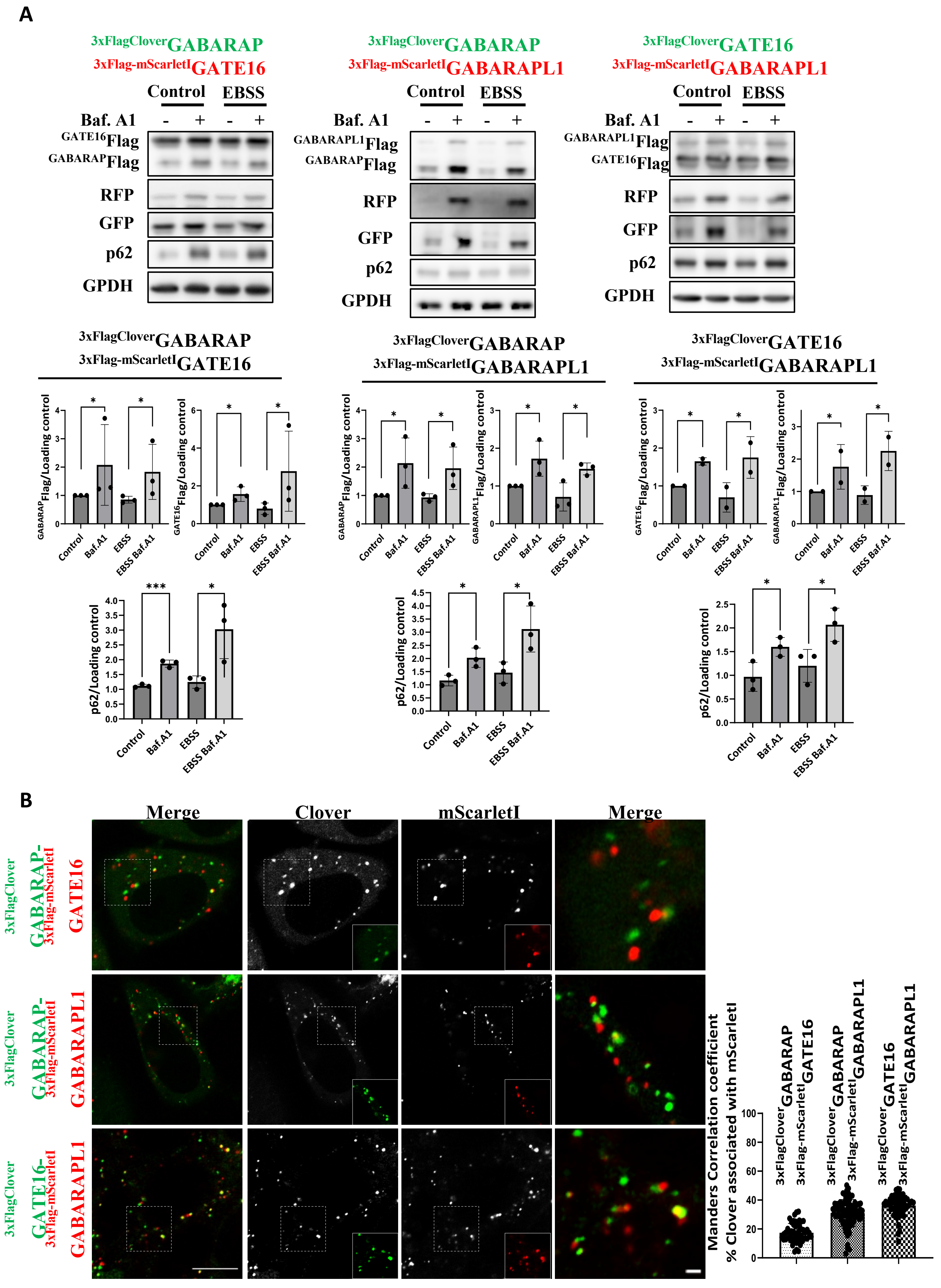


**Figure 3. Figure Supplement 1.**

### Figure 3 Supplement figure 2

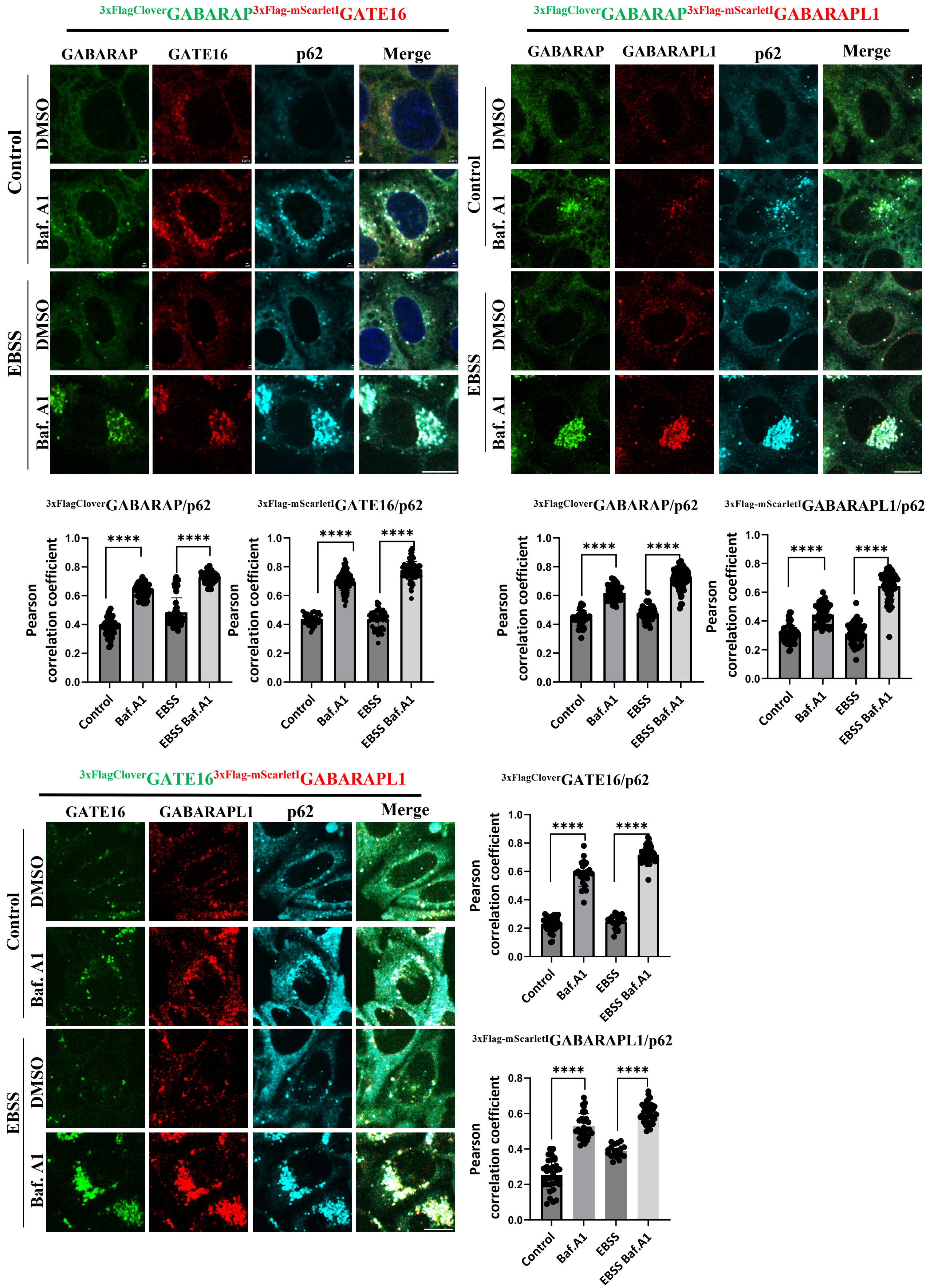


**Figure 3. Figure Supplement 2.**

### Figure 3 Supplement figure 3

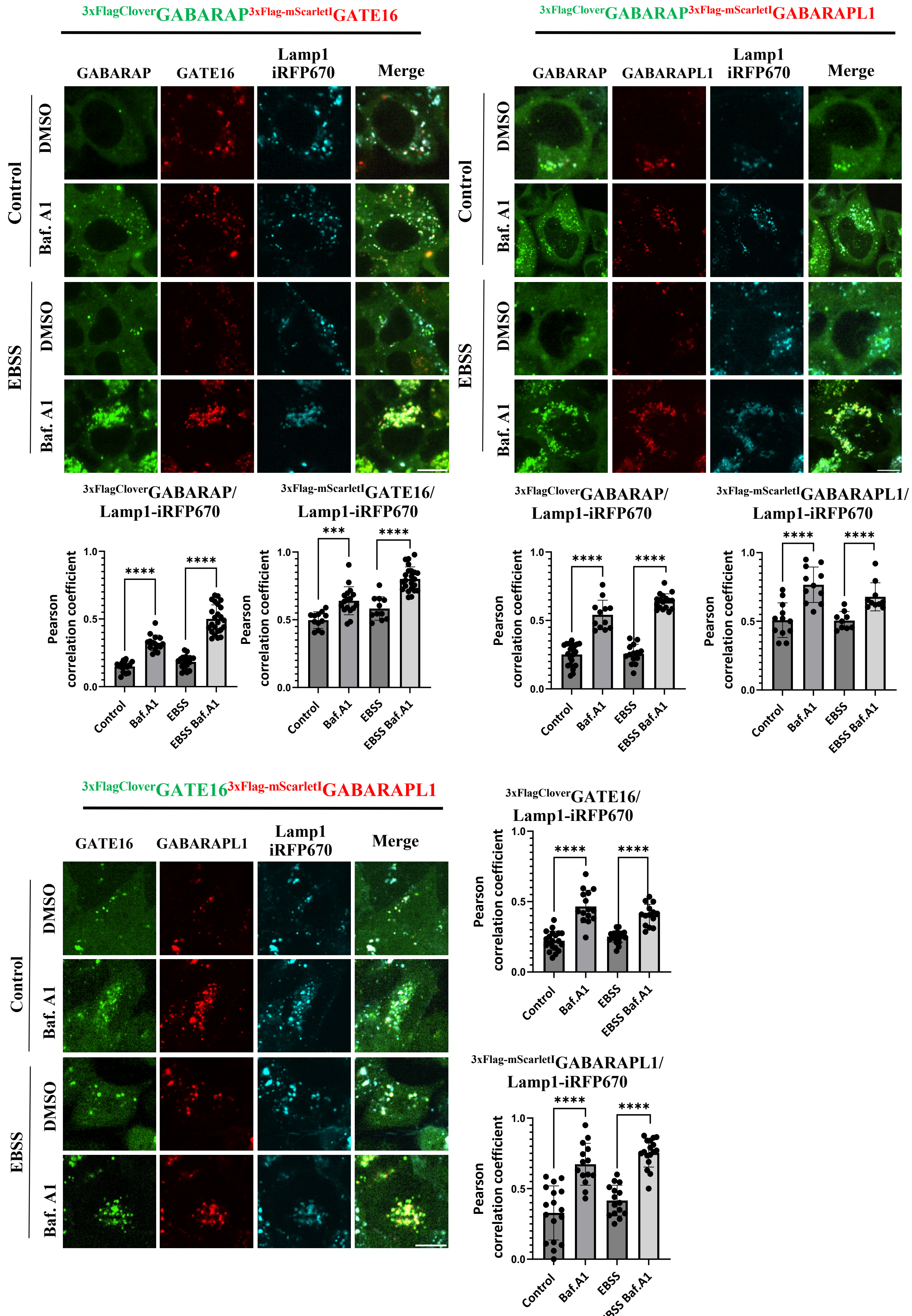


**Figure 3. Figure Supplement 3.**
